# Does stimulus order affect central tendency and serial dependence in vestibular path integration?

**DOI:** 10.1101/2025.06.20.660787

**Authors:** Sophie C.M.J. Willemsen, Leonie Oostwoud Wijdenes, Robert J. van Beers, Mathieu Koppen, W. Pieter Medendorp

**Author notes:** Corresponding author (SCMJW).

## Abstract

The reproduction of a perceived stimulus, such as a distance or a duration, is often influenced by two biases. Central tendency indicates that reproductions are biased toward the mean of the stimulus distribution. Serial dependence reflects that the reproduction of the current stimulus is influenced by the previous stimulus. Although these biases are well-documented, their origins remain to be determined. Studies on duration reproduction suggest that autocorrelation within a stimulus sequence may play a role. In this study, we explored whether the level of autocorrelation in a stimulus sequence affects central tendency and serial dependence in vestibular path integration. Participants (*n* = 24) performed a vestibular distance reproduction task in total darkness by actively replicating a passively moved stimulus distance with a linear motion platform. We compared two conditions: a high-autocorrelation condition, where stimulus distances followed a random walk, and a no-autocorrelation condition, where the same distances were presented in a randomly shuffled order. We quantified both biases using two approaches: separate simple linear regressions and a joint multiple linear regression model that accounts for the autocorrelation in the stimulus sequence. Simple linear regressions revealed that central tendency was weaker and serial dependence reversed in the high-autocorrelation condition compared to the no-autocorrelation condition. However, these differences were no longer observed in the multiple linear regression analysis, indicating that these biases were independent of the specific stimulus sequence protocol. We conclude that these perceptual biases in vestibular path integration persist regardless of stimulus autocorrelation, suggesting that they reflect robust strategies of the brain to process vestibular information in self-motion perception.

**Author summary:** How are we able to successfully navigate our surroundings? An essential part of navigation is distance estimation based on self-motion signals. We previously found that distance reproductions based on vestibular self-motion signals were affected by stimulus history. Reproductions showed a central tendency toward the mean of the stimulus distribution and an attractive serial dependence toward the immediately preceding stimulus distance. The stimulus distances were presented in a low-autocorrelation, randomized order. Here we ask whether reproductions show the same central tendency and serial dependence when consecutive stimulus distances are similar (i.e., in a high-autocorrelation, random-walk order). Participants performed a distance reproduction task in the dark: a linear motion platform first passively moved the participant over a stimulus distance, after which they actively reproduced this distance by steering the platform back to the estimated start position. We found that the reproductions showed similar central tendency and attractive serial dependence in both a no- and high-autocorrelation condition, but only if the analysis accounted for the covariation of the two effects in the high-autocorrelation condition. In conclusion, our findings indicate that central tendency and serial dependence of vestibular distance reproductions are not a result of the stimulus sequence protocol, but have neurocognitive origins.

## Introduction

Two perceptual biases that are often observed in reproduction tasks are central tendency and serial dependence. Central tendency is the notion that the participant’s reproductions tend to be biased toward the mean of the underlying stimulus distribution [1]. This bias typically leads to an overestimation of smaller stimuli and an underestimation of larger stimuli [2–15]. Serial dependence reflects that reproductions depend on the stimulus presented on the preceding trial [16,17]. Most studies have identified attractive serial dependence, where the reproduction on the current trial is biased towards the stimulus on the previous trial [15,18–23]. However, other research has reported repulsive serial dependence, indicating that the reproduction of the current stimulus is biased away from the previous stimulus [14,24].

Central tendency and serial dependence have been found to affect the perception of various stimuli, such as time durations, heading directions and distances. Veridical distance perception is essential for path integration, a process in which one uses self-motion signals to continuously estimate one’s position relative to a starting point [25,26]. These signals can be derived from our sensory systems, including the visual and vestibular systems [27], as well as the motor system [28–30]. In previous work on path integration, where participants had to mainly rely on the vestibular sense, we found central tendency and attractive serial dependence effects [15]. However, what causes these perceptual biases in vestibular path integration is not yet understood.

Recently, Glasauer & Shi [31,32] have shown that the extent to which central tendency and serial dependence effects are present in duration reproduction tasks is affected by the autocorrelation in the stimulus sequence. When durations were presented randomly shuffled, without autocorrelation, reproductions showed central tendency and attractive serial dependence. However, when the same durations were presented in a random-walk sequence with high autocorrelation, central tendency nearly disappeared and serial dependence became repulsive.

The origin of the different results between protocols remains unclear. Are these differences caused by participants responding differently in each condition, or are they byproducts of the different levels of stimulus autocorrelation? To address this question, it is important to note that central tendency and serial dependence are statistical concepts, usually defined as linear least-squares regression slopes. As such, their values can vary significantly based on the specific regression model employed and the selection of covariates included in the model. Central tendency is often characterized as 1 minus the regression slope of reproduced distance *r*_*t*_ on stimulus distance *s*_*t*_ [15,32] or, equivalently, as the negative of the regression slope of reproduction error *e*_*t*_ (*r*_*t*_ - *s*_*t*_) on stimulus distance *s*_*t*_ (see Fig 1A; [14]). Serial dependence has been defined as the regression slope of reproduction error *e*_*t*_ on the previous stimulus *s*_*t*−1_ (see Fig 1B; [15,32]). However, as illustrated in Fig 1C, central tendency and serial dependence are not independent if there is autocorrelation in the stimulus sequence (i.e., when *s*_*t*−1_ affects *s*_*t*_; see Appendix for more details). Similarly, there could be other dependencies that affect the central tendency and serial dependence coefficients (for instance, a potential effect of *s*_*t*−1_ on *e*_*t*_ through *e*_*t*−1_; see Fig 4).

**Fig 1.**
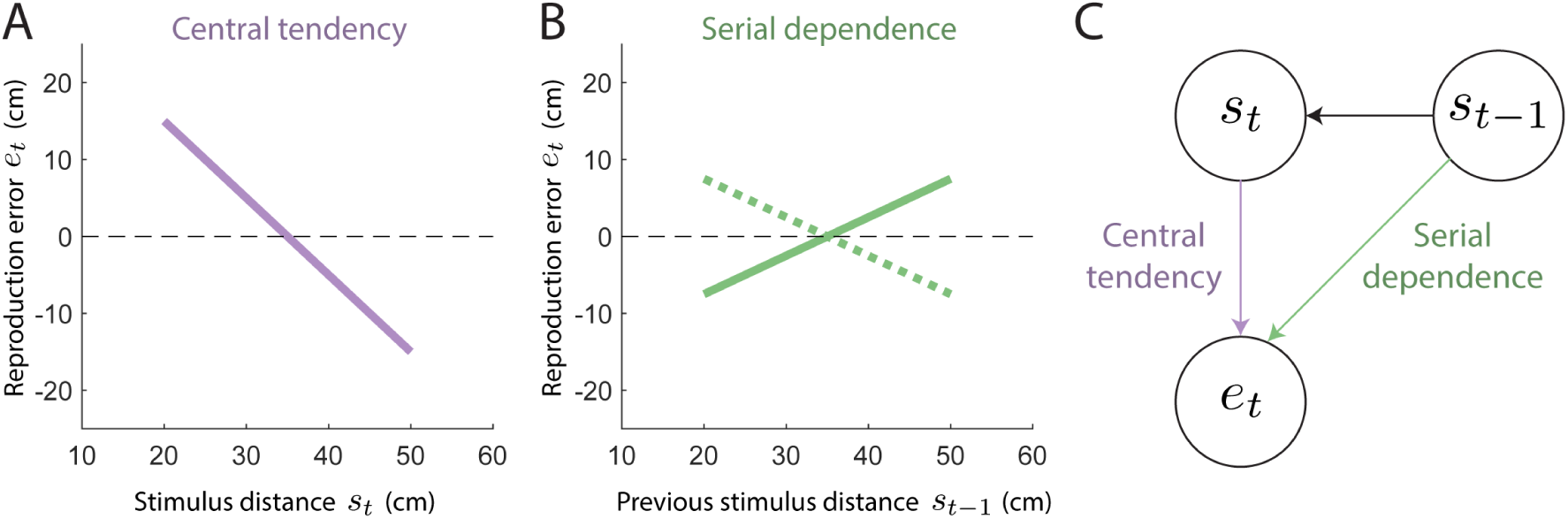
Central tendency and serial dependence. *A*: Reproduction error (reproduced distance - stimulus distance) as a function of stimulus distance. The shown line has a slope of −1, indicating a central tendency effect of 1. A regression line with a slope of 0 implies that there is no central tendency, and if also on top of the dashed line, that performance is veridical. *B*: Reproduction error against the stimulus distance on the previous trial. The solid line indicates an attractive serial dependence effect of 0.5, where the reproduction error on the current trial is generally more positive when there was a longer stimulus distance on the previous trial. The dotted line indicates a repulsive serial dependence effect of −0.5, where the reproduction error on the current trial is generally more negative when there was a longer stimulus distance on the previous trial. A regression line with a slope of 0 implies that there is no serial dependence. *C*: Central tendency (the effect of the stimulus distance on the current trial *s*_*t*_ on the reproduction error on the current trial *e*_*t*_) and serial dependence (the effect of the stimulus distance on the previous trial *s*_*t*−1_ on the reproduction error on the current trial *e*_*t*_) are not independent if there is autocorrelation in the stimulus sequence (i.e., when *s*_*t*−1_ affects *s*_*t*_).

Here, we use causal graphs and the *d*-separation criterion [33], to disentangle central tendency and serial dependence in vestibular path integration under conditions with and without stimulus sequence autocorrelation. Specifically, we ask which part of the differences in central tendency and serial dependence between the autocorrelation conditions can be attributed to a statistical explanation and which part requires an explanation in terms of different stimulus processing in the brain.

## Methods

### Participants

Twenty-five participants, naive to the purpose of the study, took part in the experiment. All participants had normal or corrected-to-normal vision, no hearing impairments and no history of motion sickness. The study was approved by the ethics committee of the Faculty of Social Sciences at Radboud University Nijmegen and all participants gave written informed consent prior to the start of the experiment. Each participant completed a single experimental session of ∼90 minutes and was compensated with course credits or €22.50. Although 24 participants were required for complete counterbalancing, one participant was excluded due to misunderstanding the task and producing reproduction movements in the wrong direction. This participant was therefore replaced by collecting data from an additional participant, resulting in a dataset of 24 participants (19 women, 4 men, 1 non-binary person, aged 17-26 yr).

### Setup

Participants were seated in a chair mounted on top of a linear motion platform, called a vestibular sled, that could be moved passively by the experimenter or actively by the participant using a steering wheel (see Fig 2). The sled was powered by a linear motor (TB15N; Tecnotion, Almelo, The Netherlands) and controlled by a servo drive (Kollmorgen S700; Danaher, Washington, DC), allowing it to move along the participant’s interaural axis on a 93-cm-long track. The steering wheel (G27 Racing Wheel; Logitech, Lausanne, Switzerland) was attached to a table at chest level in front of the participant and had a rotation range of −450° to +450° with a resolution of 0.0549°. Throughout the experiment, the mapping between the steering wheel angle and the sled’s linear velocity was set at 1 cm/s per degree. The task was performed in total darkness without any visual stimuli. Instruction messages prior to the task, as well as occasional messages throughout the experiment (e.g., to indicate breaks) were shown on an OLED screen (OLED77C3PUA; LG, Seoul, South Korea) placed in front of the sled. Participants wore in-ear headphones with active noise cancellation (QuietComfort 20; Bose, Framingham, MA) that played white noise to mask sound from the sled’s motion, alternated by single-tone beeps to signal the different stages of each trial. In addition to the in-ear headphones, participants wore over-ear headphones with active noise cancellation (WH-1000XM5; Sony, Tokyo, Japan) to further block out the sound produced by the sled. The participant’s head was fixated using cups placed against the top of the head. The participant also wore a five-point seat belt and could press one of the emergency buttons at the side of the chair to stop the sled at any time during the experiment. The experiment code was written in Python (v.3.10; Python Software Foundation).

**Fig 2.**
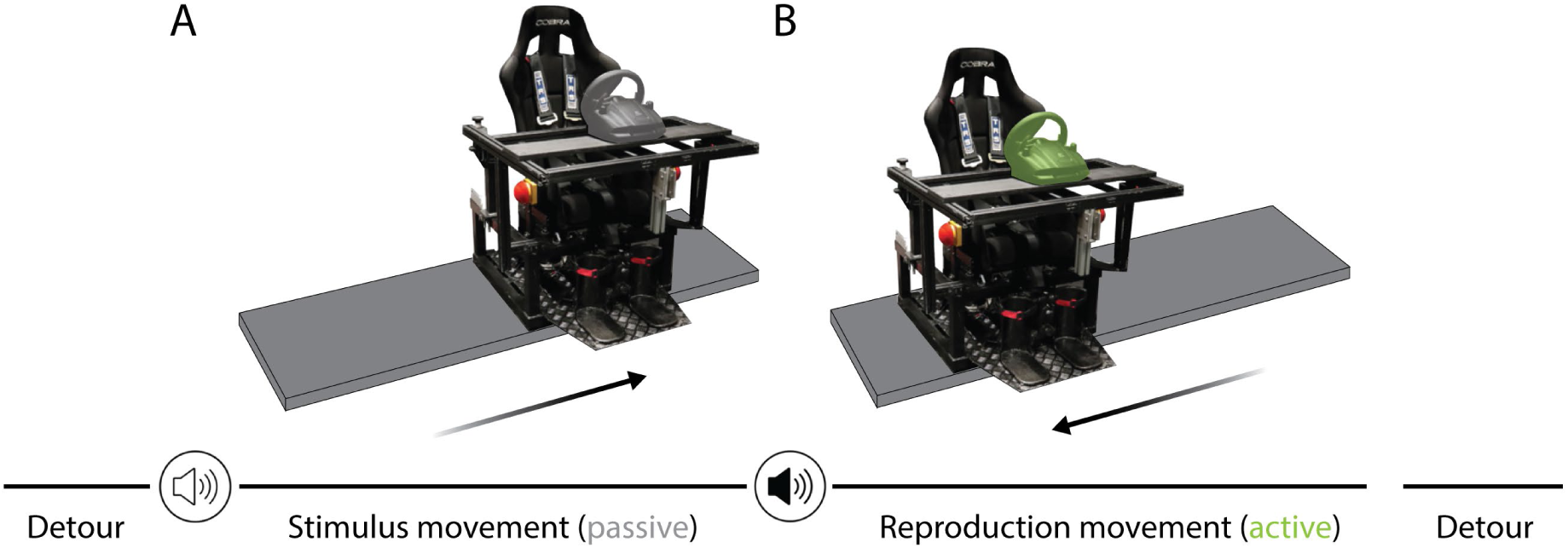
Vestibular distance reproduction task. *A*: The participant was seated on a vestibular sled, consisting of a chair placed on top of a linear motion platform. On every trial, a low-tone beep alerted the participant to the upcoming passive movement that would move them by an unknown stimulus distance. *B*: Afterwards, the second, high-tone beep prompted the participant to use the steering wheel and reproduce the stimulus distance by steering the sled back in the opposite direction. Trials were separated by two random detour movements that returned the sled to the start position.

### Reproduction task

While seated on the vestibular sled, participants performed a distance reproduction task. During the stimulus movement, the sled passively moved the participant a predefined distance (see Fig 2A). This was succeeded by the reproduction movement, during which the participant actively tried to replicate the passively moved distance by steering the sled into the opposite direction (see Fig 2B). In other words, the participant aimed to return to the start position of the stimulus movement.

In each trial, a low-tone beep (347 ms) indicated the upcoming stimulus movement. The duration of the stimulus movement varied randomly between 1.3 s and 1.6 s. We defined the lower bound such that all stimulus movements had a peak absolute acceleration below 980 cm/s^2^ and a peak speed below 100 cm/s. The upper bound resulted in the shortest stimulus movement to have a peak absolute acceleration of ∼38 cm/s^2^ and a peak speed of ∼20 cm/s, which well surpassed the vestibular thresholds [34]. For each participant, the stimulus movements were consistently in one direction, with the leftward and rightward directions counterbalanced across participants. Per participant, all stimulus movements started from the same start position, which was on the right side of the track for leftward stimulus movements and on the left side of the track for rightward stimulus movements, ensuring enough space on the track for all potential stimulus movements. The start position was determined for every participant individually depending on their largest stimulus distance. In the case of leftward stimulus movements, the start position was determined by adding the largest stimulus distance to the leftmost position on the sled track plus an additional small margin of 4 cm. For rightward stimulus movements, the start position was computed by subtracting the largest stimulus distance and the margin from the rightmost position on the track.

The stimulus movement was followed by a random waiting time between 0.5 s and 1 s, after which a high-tone beep (110 ms) cued the start of the reproduction movement. If the participant rotated the steering wheel before the beep, the trial was aborted. Participants were instructed to make one smooth reproduction movement (without steering back or resuming steering after stopping) and were free to choose the duration of the movement. The sled could be steered up to a maximum speed of 100 cm/s and could be stopped by returning the steering wheel back to the upright position. The movement was terminated when the speed fell below 2 cm/s. The sled also stopped moving when the speed fell below 6 cm/s while the steering angle remained unchanged for 100 ms or the steering changed direction (mean ± SD across participants: 71 ± 59 trials out of a total of 260 trials). This second stopping criterion was added to prevent the case where the participant intended to stop the movement but did not fully return the steering wheel to the upright position. When one of these stopping criteria was met, the sled would not stop abruptly but would decelerate in 1 s to a speed of 0 cm/s. The sled also stopped moving when the end of the sled track was reached (mean ± SD: 2 ± 2 trials).

Participants received no feedback about their reproduction performance during the experiment (except during the training block, see below). To prevent the participant from obtaining implicit feedback about their reproduced distance, the sled was brought back to the start position for the next stimulus movement through two random detour movements. The first detour relocated the sled to a random position within ±30 cm from the middle of the track with a random duration between 1.8 s and 2.3 s. The second detour moved the sled to the start position in 1.3 s. All detour and stimulus movements followed a minimum-jerk profile.

### Paradigm

To study how the amount of autocorrelation between the stimulus distances across trials affects central tendency and serial dependence biases in vestibular path integration, we created two experimental conditions per participant presenting the same stimulus distances with different stimulus orders. In the high-autocorrelation condition, stimulus distances followed a random walk while in the no-autocorrelation condition, the same distances were randomly shuffled (see Fig 3).

**Fig 3.**
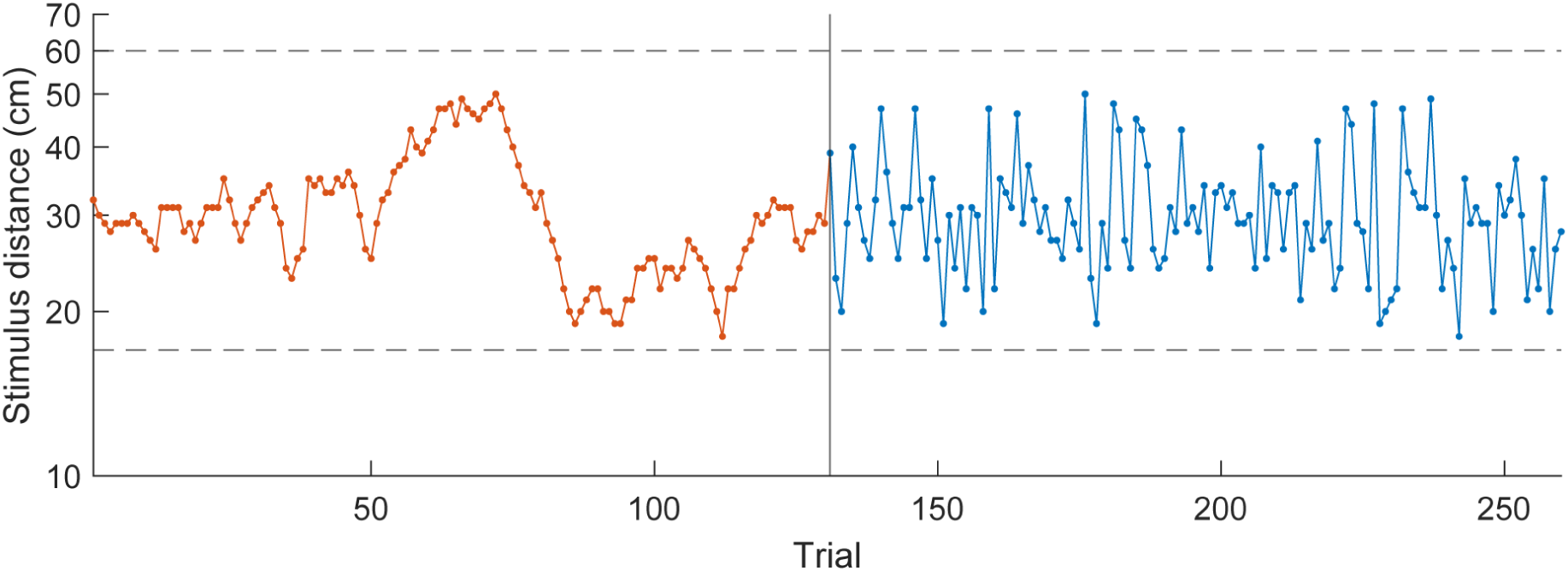
Example sequence of stimulus distances. Stimulus distances throughout the entire experimental session for a participant starting with the high-autocorrelation condition. During the first 130 test trials, the stimulus distances followed a random walk on logarithmic scale (orange). In the second half of the experiment, the same distances were presented in a randomly shuffled order (blue). The dashed lines indicate the minimum and maximum possible stimulus distance.

**Fig 4.**
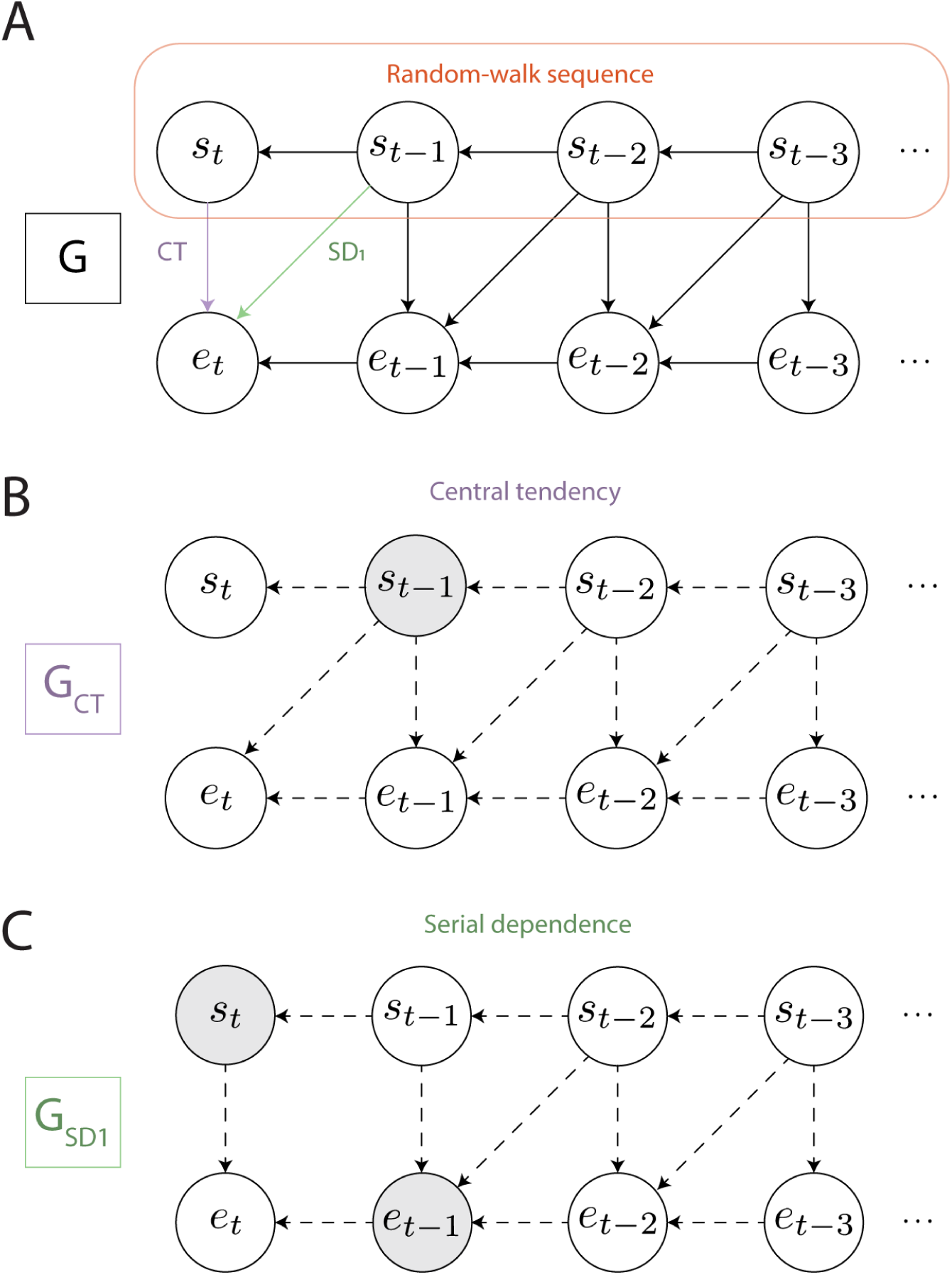
Causal modeling. *A*: Causal diagram *G* representing the assumed causal relationships between the stimulus distances (*s*) and reproduction errors (*e*) across trials (*t*) in the high-autocorrelation condition. Variables are presented by nodes and possible causal relationships between the variables by directed edges. A path between two variables denotes a set of edges that connects the two variables (irrespective of the direction of the edges). The upper row of nodes represents the random-walk sequence, in which the previous stimulus affects the current stimulus. The edge *CT* between the current stimulus *s*_*t*_ and the current reproduction error *e*_*t*_ reflects the possible central tendency effect. Similarly, the edge *SD*_1_ between the previous stimulus *s*_*t*−1_ and the current reproduction error *e*_*t*_ represents the possible serial dependence effect at lag 1. *B*: Application of the single-door criterion to determine which variables to include as regressors in a multiple linear regression model such that central tendency coefficient *CT* is identifiable. Graph *G*_*CT*_ is equal to graph *G* with edge *CT* removed. Dashed arrows indicate (parts of) blocked paths between *s*_*t*_ and *e*_*t*_ and gray nodes represent the variables to add as regressors.

For each participant, we first generated 130 stimulus distances following a random walk. In line with our previous study [15], the random walk was generated on logarithmic scale such that the resulting stimulus distances were approximately normally distributed on this scale. For this transform, distances were made dimensionless by dividing by a reference distance (1 cm). On a linear scale, the distances varied between 17 cm and 60 cm and the first distance of the random-walk sequence was set to the median of this distance range on logarithmic scale, which corresponds to 31.9 cm on linear scale. To create the remainder of the sequence, 129 random shifts were drawn from a normal distribution with a mean of 0 and SD of 0.08, and these were cumulatively summed to the first distance. Across participants, the stimulus distances on logarithmic scale varied between 2.83 and 4.05, the mean of the sequence between 3.37 and 3.50, the SD of the sequence between 0.20 and 0.27 and the lag-1 autocorrelation was larger than 0.9. We computed the lag-1 autocorrelation *r*_1_ [35] using

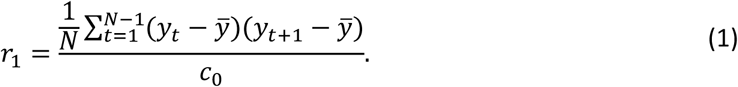

Here, the numerator is the autocovariance of the sequence which is divided by the sample variance of the sequence *c*_0_, resulting in an autocorrelation value between −1 and 1. Furthermore, *N* denotes the total number of samples in the sequence and *y^-^* the sample mean of the sequence. To create the no-autocorrelation condition, the same 130 stimulus distances were shuffled until the autocorrelation of the sequences was between −0.001 and 0.001.

Participants experienced both conditions in a single experimental session of 260 test trials (see Fig 3 for an example sequence of stimulus distances) without being informed about the presence of the two conditions. The order of the conditions was counterbalanced across participants. There was a short break (∼2 min) after every 52 trials (∼10 min) with the room lights turned on to prevent dark adaptation.

Prior to the test trials, participants completed 20 training trials to get acquainted with the task. The stimulus distances on the training trials were drawn from a uniform distribution between 17 cm and 60 cm on linear scale. The training trials were performed in darkness and differed from the test trials in two respects. During the first 10 training trials, instruction texts were displayed on the screen alongside the beeps to indicate the various trial phases. Four instruction texts were shown for each trial, preceding the first detour, the second detour, the stimulus movement and the reproduction movement, respectively. In the second half of the training trials, these instruction texts were not shown such that only the beeps indicated the different trial phases. The second difference with the test trials was that participants received feedback about their performance, displayed as the signed reproduction error in centimeters at the end of each training trial. We did not analyze the training trials.

### Data analysis

#### Pre-processing

We analyzed data from the test trials offline in MATLAB (v.R2019a, MathWorks). The end position of the reproduction movement was defined as the sled position at the moment when the participant moved the steering wheel upright. We chose this position as opposed to the sled position after the slow-down period, as it more accurately reflects the participant’s intended end position. Some of the recorded sled position profiles indicated that movement speed plateaued at a low but nonzero value before the slow-down period was initiated. The movement end was therefore corrected to the first time point where sled speed was < 8 cm/s (instead of the online threshold of 6 cm/s) while the steering angle remained constant for at least 100 ms or the steering direction changed. On average, the end position of 20 trials per participant were determined in this way (mean ± SD: 20 ± 15 trials). The reproduction error was computed as reproduced distance minus stimulus distance on logarithmic scale, with negative values indicating an undershoot and positive values an overshoot. We excluded trials if the participant initiated the reproduction movement too early, if reproduction movements were in the wrong direction, or if the reproduced distance was less than 1 cm (mean ± SD: 5 ± 6 trials). There was no effect of movement direction on the mean unsigned reproduction error across trials (Wilcoxon rank-sum test, *p* = 0.624, rank-biserial correlation = 0.13), so participants were analyzed as a single group, disregarding this factor.

#### Central tendency and serial dependence computation

Fig 4A illustrates a causal diagram *G* [33] of the high-autocorrelation condition. Variables are represented as nodes and possible causal relationships between the variables as directed edges. A path between two variables consists of a set of edges that connects the two variables (irrespective of the direction of the edges). In this diagram, the *s*-nodes represent the stimulus distances and the *e*-nodes the reproduction errors at different trials *t*. As the stimulus distances in this condition are presented in a random-walk sequence, we know that the current stimulus *s*_*t*_ depends on the previous stimulus *s*_*t*−1_ which in turn depends on *s*_*t*−2_ and so forth (the top row in Fig 4A). Furthermore, the current reproduction error may be affected by the current and previous stimulus distances (the vertical and diagonal edges in Fig 4A) and the previous reproduction error (the bottom row in Fig 4A).

The edge between the current stimulus *s*_*t*_ and the current reproduction error *e*_*t*_ represents the central tendency effect, whose coefficient *CT* we want to estimate. A negative coefficient suggests a central tendency effect, where the longer the stimulus distance is, the more it is underestimated (i.e., the more negative the reproduction error becomes). Finding a coefficient of 0 implies that there is no central tendency effect (i.e., the reproduction error is constant across stimulus distances) and a positive coefficient implies that there is anti-central tendency in the reproductions.

Similarly, the edge from the stimulus distance on the previous trial *s*_*t*−1_ to the reproduction error on the current trial *e*_*t*_ captures the serial dependence effect at lag 1. Here, we express serial dependence as the dependence of the current error on the previous stimulus distance (‘absolute’ serial dependence; e.g., Holland & Lockhead [16]) instead of the dependence of the current error on the difference between the previous stimulus and the current stimulus (‘relative’ serial dependence; e.g., Fischer & Whitney [18]). The latter metric can erroneously result in a serial dependence effect if stimuli are defined on an open scale (such as distances or durations) and the reproductions are constant across stimuli (see Appendix A in Glasauer & Shi [32]). A positive (attractive) serial dependence coefficient *SD_1_* indicates that if the participant experienced a longer stimulus distance on the previous trial, they tend to show a larger overestimation (i.e., a more positive reproduction error) on the current trial. A coefficient of 0 implies that there is no serial dependence and a negative (repulsive) coefficient reflects that a longer stimulus distance on the previous trial tends to be followed by a larger underestimation (i.e., a more negative reproduction error) on the current trial.

As becomes apparent from the graph in Fig 4A, beside the direct path *CT* there are indirect paths through which *s*_*t*_ can affect *e*_*t*_. For example, there exists an indirect path from *s*_*t*_ to *e*_*t*_ via common cause *s*_*t*−1_. In order to accurately estimate the coefficient of the direct path *CT*, this indirect path should be ‘blocked’ by adding variable *s*_*t*−1_ to the adjustment set *Z*, i.e., by adding this variable as a regressor to the multiple linear regression model (*e*_*t*_ = *CT* ⋅ *s*_*t*_ + β ⋅ *s*_*t*−1_ + ε). More generally, all indirect paths that connect *s*_*t*_ and *e*_*t*_ should be blocked, in which case *s*_*t*_ is said to be *d*-separated from *e*_*t*_. The coefficient *CT* is said to be identifiable when there exists an adjustment set *Z* that *d*-separates *s*_*t*_ from *e*_*t*_ and when *Z* contains no descendants of *e*_*t*_ (Theorem 5.3.1., the single-door criterion for direct effects [33]). If these conditions are not satisfied, this may lead to a biased estimate of *CT*.

Beside the direct path *CT*, we can see that all indirect paths between *s*_*t*_ and *e*_*t*_ contain *s*_*t*−1_, so by adding this variable to the adjustment set, all indirect paths between *s*_*t*_ and *e*_*t*_ are blocked (see Fig 4B). Similarly, to *d*-separate *s*_*t*−1_ and *e*_*t*_ (beside the direct path *SD_1_*), *s*_*t*_ and *e*_*t*−1_ should be adjusted for, blocking all indirect paths between *s*_*t*−1_ and *e*_*t*_ (see Fig 4C). Thus, to estimate the direct *CT* effect, the regression of *e*_*t*_ on *s*_*t*_ also has to include the regressor *s*_*t*−1_ and to estimate the direct *SD_1_* effect, the regression of *e*_*t*_ on *s*_*t*−1_ also has to include the regressors *s*_*t*_ and *e*_*t*−1_. As the latter regression model contains the first (and adding the *e*_*t*−1_ regressor to the *CT* regression model does not open up paths between *s*_*t*_ and *e*_*t*_), we combine the two multiple linear regression models. This results in one model that can be used to estimate both the central tendency effect *CT* and the serial dependence effect *SD_1_*:

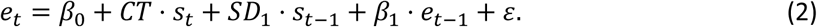

A similar causal graph can be drawn for the no-autocorrelation condition, but without edges between the stimulus distances. From this graph follows that to estimate *CT*, no variables have to be adjusted for and to estimate *SD_1_*, *e*_*t*−1_ should be adjusted for. The same regression model as above can also be used to estimate the central tendency and serial dependence effects in the no-autocorrelation condition, because indirect paths between *s*_*t*_ and *e*_*t*_ remain blocked when also adjusting for *s*_*t*−1_ and *e*_*t*−1_, and indirect paths between *s*_*t*−1_ and *e*_*t*_ remain blocked when also adjusting for *s*_*t*_.

By adding *s*_*t*−1_ as a regressor, all biasing paths between *s*_*t*_ and *e*_*t*_ are blocked and *CT* can be estimated. *C*: Application of the single-door criterion to serial dependence coefficient *SD1*. By adding *s*_*t*_ and *e*_*t*−1_ as regressors, all biasing paths between *s*_*t*−1_ and *e*_*t*_ are blocked and *SD1* becomes identifiable.

To compare the central tendency and serial dependence across the autocorrelation conditions, we fitted this model to the data of each participant and each condition separately, on logarithmic scale. Partial regression plots of the current reproduction error on the current stimulus distance, and of the current reproduction error on the previous stimulus distance are used to visualize the central tendency and serial dependence effects, respectively. These plots were created using the MATLAB function *plotAdded* and illustrate the effect of one regressor on the response variable while keeping the other regressors constant. The slope of the fitted line corresponds to the fitted partial regression coefficient (*CT* and *SD_1_*, respectively).

To illustrate how accounting for the biasing paths affects the central tendency and serial dependence coefficients, we also computed the same coefficients by fitting two separate simple linear regressions to the data of each participant and condition. The models used to estimate the central tendency effect *CT* and the serial dependence effect *SD_1_* were *e*_*t*_ = β_0_ + *CT* ⋅ *s*_*t*_ + ε and *e*_*t*_ = β_0_ + *SD*_1_ ⋅ *s*_*t*−1_ + ε, respectively. The Appendix provides a comparison of the different central tendency and serial dependence metrics based on simulated data.

#### Statistical tests

To further analyze the central tendency and serial dependence coefficients, we used the following statistical tests. We first tested whether there was an effect of condition (no/high-autocorrelation) on the central tendency/serial dependence coefficients with paired-samples *t*-tests. One-sample *t*-tests were used to assess whether the central tendency/serial dependence coefficients significantly differed from 0. Cohen’s *d* [36] and 95% confidence intervals are reported.

## Results

Participants performed a vestibular distance reproduction task in the dark where they actively reproduced a stimulus movement that they had passively experienced. To manipulate the level of autocorrelation of the stimulus distances, we established a high-autocorrelation condition characterized by a random walk of stimulaai, alongside a no-autocorrelation condition where the same stimuli were randomly shuffled. We examined the central tendency and serial dependence effects on the reproductions in these conditions.

### Central tendency

Fig 5, A and B, present the simple linear regressions of reproduction error versus stimulus distance on the current trial, without adjusting for covariates, for a single participant in the no- and high-autocorrelation conditions, respectively. The slope of the fitted regression line corresponds to the central tendency coefficient *CT*. In both conditions, *CT* is negative indicating central tendency, with the high-autocorrelation condition showing less central tendency than the no-autocorrelation condition. For comparison, Fig 5, C and D, show the partial regression plots of the same participant based on the multiple linear regression model (see Equation 2), with adjustment for covariates. These adjusted values indicate that the effective variance in the stimulus distances is lower in the high-autocorrelation than the no-autocorrelation condition. In contrast to the analysis presented in Fig 5, A and B, the partial regression coefficients suggest that central tendency remains fairly consistent across conditions.

**Fig 5.**
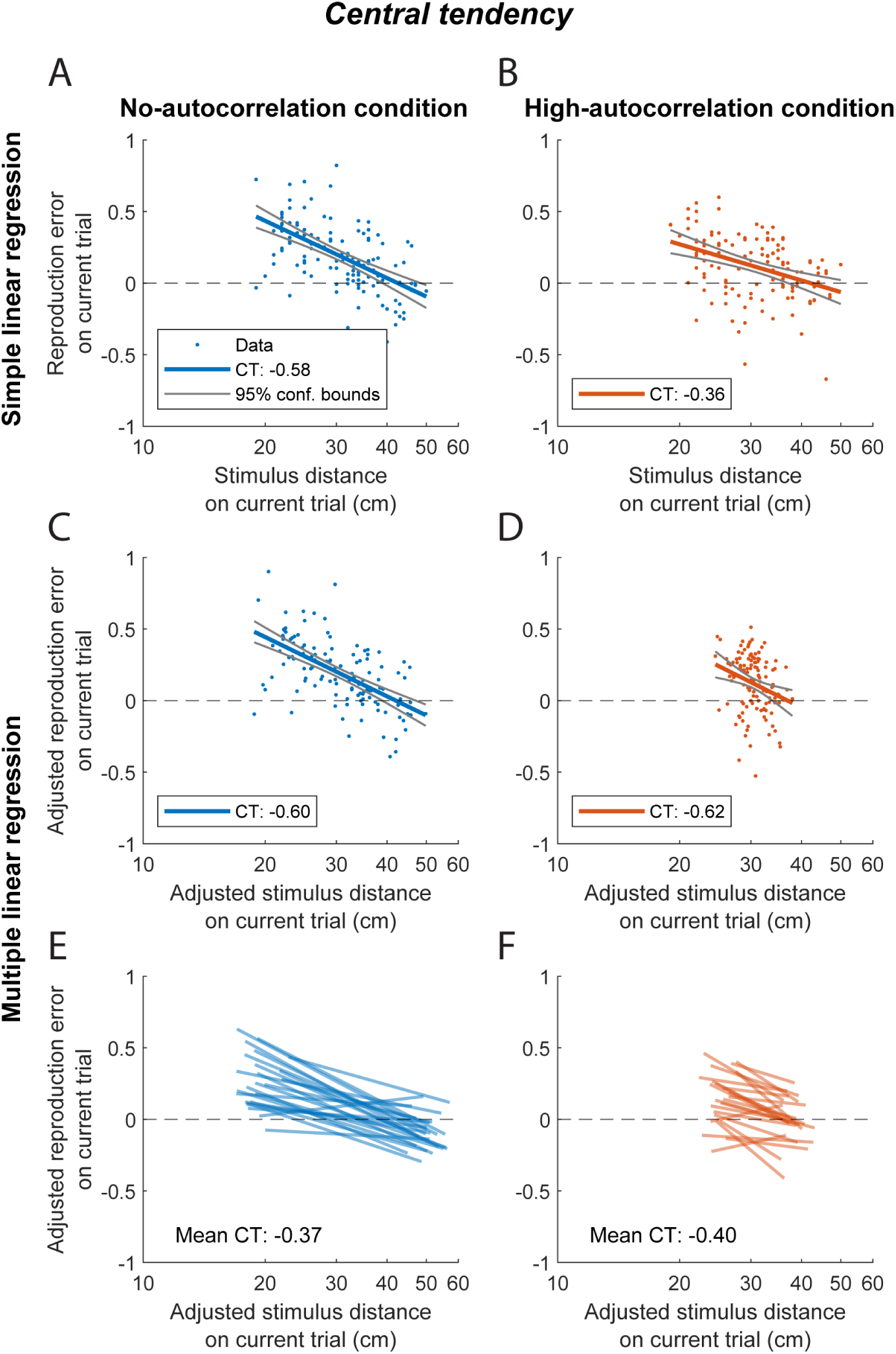
Central tendency regressions. Regression plots of the reproduction error on the current trial as a function of stimulus distance on the current trial in the no-autocorrelation (blue) and high-autocorrelation (orange) conditions on logarithmic scale for an individual participant (*A*-*D*) and all participants (*E*-*F*). Regression lines illustrate the central tendency, with the regression slope corresponding to the regression coefficient *CT*. *A*-*B*: Simple linear regression lines, with the *CT* value reported in the key. *C*-*D*: Partial regression lines based on the multiple linear regression model. *E-F*: Partial regression lines with the mean fitted *CT* coefficient across participants indicated.

Fig 5, E and F, illustrate the regression lines for all participants, as estimated by the multiple linear regression model, which reveal no significant difference in central tendency between the conditions (paired-samples *t*-test: *p* = 0.550, Cohen’s *d* = 0.12, 95% CI = [−0.07, 0.14]). As visualized in Fig 6A, the *CT* values exhibit considerable variability between participants. Yet, they are on average negative across conditions (M = −0.38, SD = 0.29), indicating a substantial level of central tendency (one-sample *t*-test: *p* < 0.001, Cohen’s *d* = 1.35, 95% CI = [−0.46, −0.31]).

**Fig 6.**
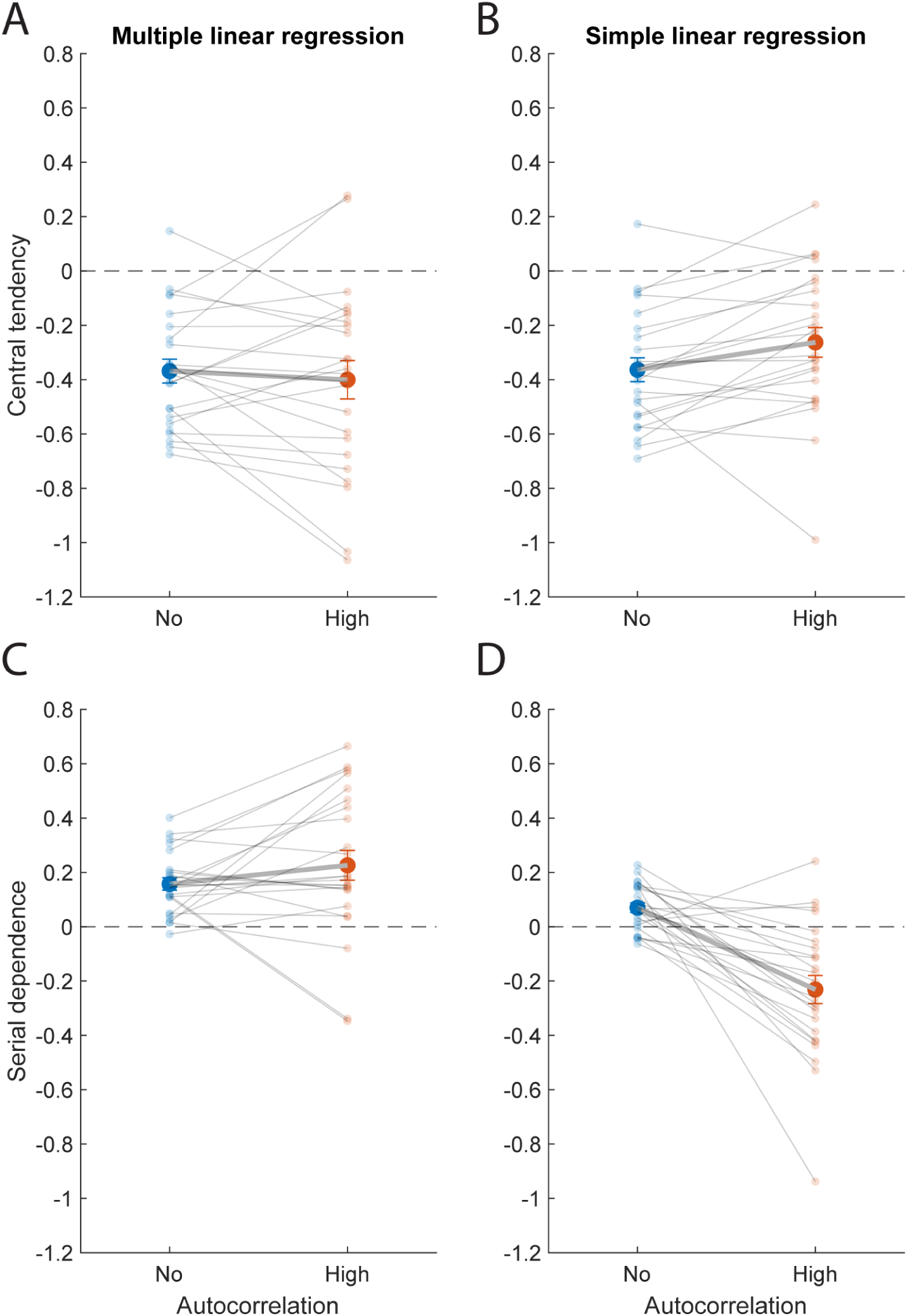
Central tendency and serial dependence regression coefficients. Central tendency (*A*, *B*) and serial dependence (*C*, *D*) regression coefficients in the no-autocorrelation (blue) and high-autocorrelation (orange) conditions. Panels *A* and *C* show the partial regression coefficients computed with the multiple linear regression model, and panels *B* and *D* show the regression coefficients computed with the two separate simple linear regression models. Bold data points and error bars represent the mean ± SE across participants. Transparent data points and their connecting lines show individual participants.

For comparison, Fig 6B presents the *CT* values, as calculated with the simple linear regression. In both conditions, the mean *CT* coefficient across participants is significantly smaller than zero (no-autocorrelation: M = −0.36, SD = 0.21, *p* < 0.001, Cohen’s *d* = 1.70, 95% CI = [−0.45, −0.28], high-autocorrelation: M = −0.26, SD = 0.27, *p* < 0.001, Cohen’s *d* = 0.98, 95% CI = [−0.37, −0.16]). More strikingly, the average *CT* values differed significantly between the two conditions (paired-samples *t*-test: *p* = 0.016, Cohen’s *d* = 0.53, 95% CI = [−0.18, −0.03]), demonstrating that not accounting for the autocorrelation in the stimulus sequence can result in different central tendency coefficients.

### Serial dependence

Fig 7, A and B, show simple regression plots of the same exemplary participant as in Fig 5, but now with reproduction error on the current trial plotted against stimulus distance on the previous trial. The regression line illustrates the serial dependence, of which the slope corresponds to the fitted regression coefficient *SD*_1_. In the no-autocorrelation condition, the positive *SD*_1_ indicates that there is attractive serial dependence, whereas this coefficient is negative in the high-autocorrelation condition, representing repulsive serial dependence. Fig 7, C and D, display regression plots of the same data set adjusted for the other regressors in the multiple linear regression model (see Equation 2). Compared to the simple regression analysis, *SD*_1_ remains positive in the no-autocorrelation, but shifts from negative to positive in the high-autocorrelation condition.

**Fig 7.**
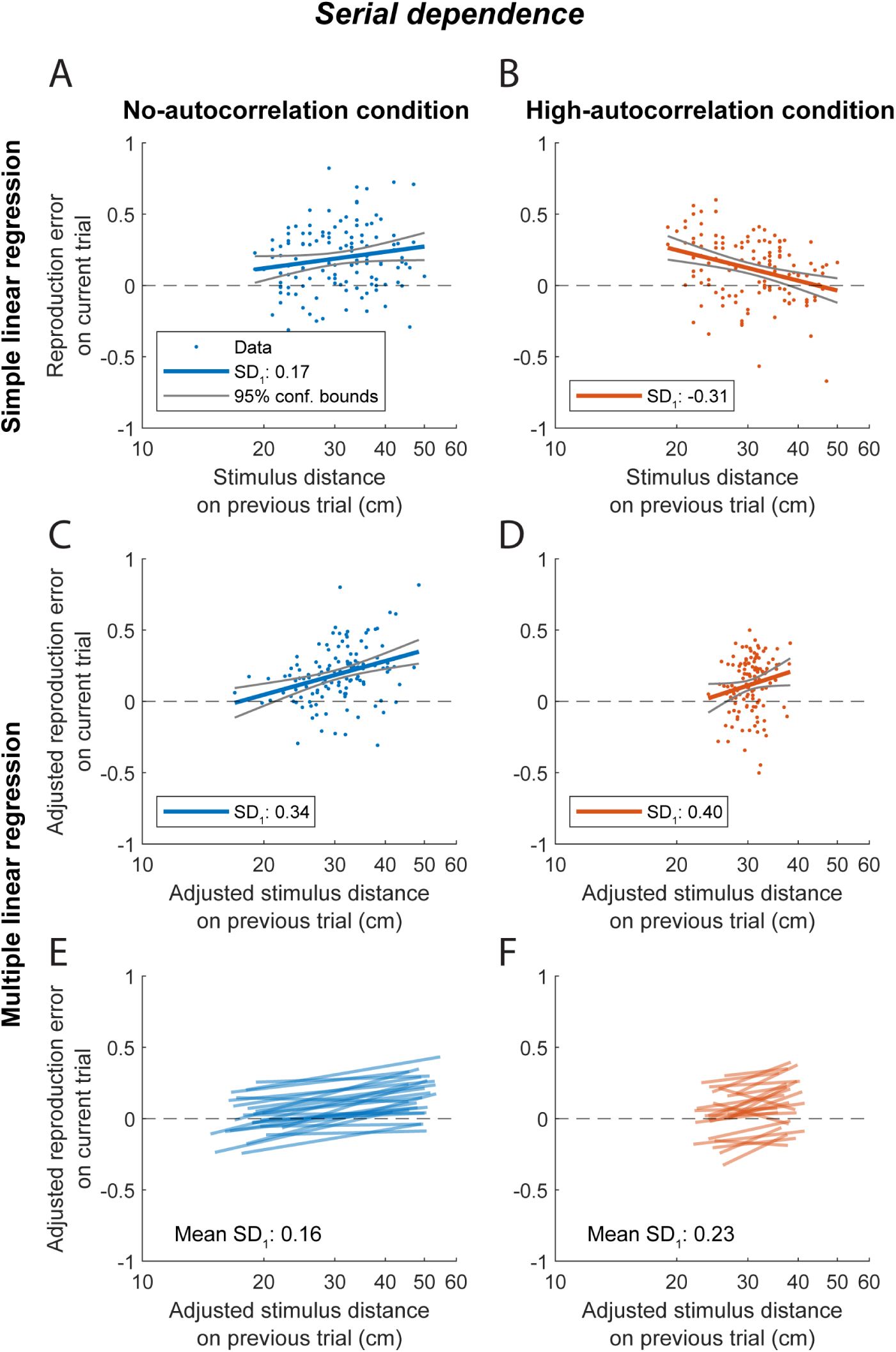
Serial dependence regressions. Regression plots of the reproduction error on the current trial as a function of stimulus distance on the previous trial in the no-autocorrelation (blue) and high-autocorrelation (orange) conditions on logarithmic scale. Regression lines illustrate the serial dependence, with the regression slope corresponding to the regression coefficient *SD1*. The figure is in the same format as Fig 5, with the same individual participant.

Fig 7, E and F, display the serial dependence lines for all participants, as determined by the multiple linear regression model. A paired-samples *t*-test indicated no significant difference between the average *SD*_1_ coefficients of the no- and high-autocorrelation conditions (*p* = 0.180, Cohen’s *d* = 0.28, 95% CI = [−0.17, 0.03]). Despite substantial intersubject variability (see Fig 6C), *SD*_1_ was on average positive across conditions (M = 0.19, SD = 0.21), suggesting attractive serial dependence (one-sample *t*-test: *p* < 0.001, Cohen’s *d* = 0.93, 95% CI = [0.14, 0.25]).

For comparison, Fig 6D shows these coefficients, as determined from fitting the simple linear regression. In this case, a paired-samples *t*-test revealed a significant difference between the two conditions (*p* < 0.001, Cohen’s *d* = 1.10, 95% CI = [0.19, 0.41]), with attractive serial dependence in the no-autocorrelation condition (M = 0.07, SD = 0.08, one-sample *t*-test: *p* < 0.001, Cohen’s *d* = 0.82, 95% CI = [0.04, 0.10]) and repulsive serial dependence in the high-autocorrelation condition (M = −0.23, SD = 0.25, one-sample *t*-test: *p* < 0.001, Cohen’s *d* = 0.92, 95% CI = [−0.33, −0.13]). Again, this highlights that the two analysis methods can lead to different results and therefore different interpretations of the data.

## Discussion

In this study, we investigated the effect of the autocorrelation in the stimulus sequence on central tendency and serial dependence in vestibular path integration. Participants performed a distance reproduction task using a vestibular sled in total darkness. On each trial, the participant was passively moved over a specified stimulus distance, which they actively reproduced by steering the sled back to the start position. Each participant completed two experimental conditions during which the same stimulus distances were presented but in different orders. In the high-autocorrelation condition, the stimuli followed a random walk, whereas in the no-autocorrelation condition, the stimulus distances were randomly shuffled. Central tendency and serial dependence were assessed either by conducting two separate simple linear regressions or by employing a single multiple linear regression model. The latter approach was derived from a causal diagram (see Fig 4, cf. Pearl [33]), taking into account that the two perceptual biases may covary due to autocorrelated stimuli. We found that applying the two analytical methods to the vestibular path integration data set yielded different results regarding how autocorrelation influences both central tendency and serial dependence.

The simple linear regressions suggest that central tendency was weaker in the high-autocorrelation than in the no-autocorrelation condition. This approach also indicates that the level of stimulus autocorrelation can make the serial dependence coefficient flip sign: the high-autocorrelation condition demonstrated repulsive serial dependence, while the no-autocorrelation condition demonstrated attractive serial dependence. However, when we used multiple linear regression to jointly quantify both central tendency and serial dependence, thus accounting for their covariation as well as the effect of the previous reproduction error, we observed no significant differences in either perceptual bias between the two autocorrelation conditions. In both conditions, we found similar central tendency and attractive serial dependence effects, suggesting that these biases are independent of the specific stimulus sequence protocol that was used. Can we reconcile these different outcomes with findings from previous literature?

Our simple linear regression results align with the findings of Glasauer & Shi [32], who reported that both central tendency and serial dependence in reproduced durations, estimated using separate simple linear regressions, depended on the sequence of the presented stimuli. The multiple linear regression coefficients of this study are consistent with the central tendency and serial dependence biases found in our previous study on vestibular path integration [15]. In this earlier study, stimulus distances were randomly sampled from either a short- or long-distance distribution and presented in a mixed or blocked order, with stimulus autocorrelations (per distance and order type) that were on average close to 0 across participants (mean ± SD: −0.03 ± 0.13). The reproduced distances showed a similar amount of central tendency and attractive serial dependence as in the present study, for both distance types (short/long) and presentation contexts (mixed/blocked).

The novelty of the present study is that we found central tendency and serial dependence in vestibular path integration to be independent of stimulus autocorrelation, if these biases are estimated by a multiple linear regression model that accounts for their covariation. Thus, the differences in central tendency and serial dependence identified through the separate simple linear regressions are due to the different levels of autocorrelation that were not accounted for in the regressions, rather than due to differences in brain processing across the two conditions. As shown in the Appendix, separately estimating the biases in simulated reproductions that show central tendency but no serial dependence, can falsely result in a repulsive serial dependence coefficient when stimuli are autocorrelated. The autocorrelation in the stimuli makes that a short stimulus is likely to follow another short stimulus. If we tend to overestimate short stimuli irrespective of the previous stimulus (the central tendency effect), this will also show up as repulsive serial dependence, i.e., an overestimation that occurs if the previous stimulus was short.

Here, we show that central tendency and serial dependence in vestibular path integration persist regardless of stimulus autocorrelation, which suggests that they reflect robust neural processes that affect the estimation of self-motion, even when the stimulus changes predictably over time. Specifically, we found that reproductions showed central tendency: shorter stimulus distances were generally overestimated, while longer distances tended to be underestimated. This pattern aligns with previous findings in distance and heading perception, where central tendency has been consistently reported [2–6,8,9,11,14,37,38]. Furthermore, the reproduction errors showed attractive serial dependence, which indicates that self-motion perception of participants is also biased toward the stimulus distance of the immediately preceding trial. Attractive serial dependence effects have been widely reported in the perception literature [18–23]. While attractive serial dependence in vestibular path integration may help to stabilize self-motion perception from trial to trial, it would reduce sensitivity to small changes between trials [14,22].

To computationally understand the underlying neurocognitive processes, numerous studies have adopted a Bayesian framework to explain central tendency and serial dependence. In this approach, the brain is thought to encode information about previous stimuli as a prior distribution, which is optimally combined with the sensory likelihood, using Bayes’ rule [9,11,38–40]. It can be shown that if the prior and likelihood are modeled as Gaussian distributions, their combination will result in a posterior distribution with a lower variance, reflecting more precise but potentially biased estimates. Recent research indicates that the posterior parietal cortex may play a role in these computations [41].

Within the Bayesian framework, Glasauer & Shi [32] proposed a Kalman filter model that iteratively combines the sensory measurement from the current trial with the stimulus estimate from the previous trial. It can be shown that the steady state of this model is similar to an ARX model on logarithmic scale [42]. By varying the Kalman filter’s assumptions about the estimated stimulus distribution, the authors assessed how various beliefs about the generation of stimuli in the environment could explain the central tendency and serial dependence biases. Both central tendency and serial dependence effects, in duration perception as well as in visual path integration, were well explained by a model that assumes that the stimuli are drawn from a stimulus distribution of which the mean can fluctuate across trials [32]. Additionally, this model demonstrated a reasonably good fit to the vestibular distance reproductions in our previous study, successfully capturing the central tendency effects in the data, although it was less effective in explaining the serial dependence effects [15]. As the focus of the current study was on the computation of central tendency and serial dependence across different levels of stimulus autocorrelation, evaluating the fit of the Kalman filter model to the current data set was outside the scope of this study.

As a final consideration regarding the multiple linear regression analysis, it is important to note that the causal diagram from which it is derived represents an assumed causal structure underlying the high-autocorrelation condition. If relevant variables or connections are missing, there is a risk that direct effects may be misidentified. For example, earlier stimulus distances (see *s*_*t*−2_, *s*_*t*−3_, etc. in Fig 4A) might also influence the current reproduction error. The causal graph in Fig 4A implies that *e*_*t*_ and *s*_*t*−2_ are conditionally independent given *s*_*t*−1_ and *e*_*t*−1_; an assumption that we tested using the high-autocorrelation data set. We fitted the multiple linear regression model *e*_*t*_ = β_0_ + β_1_ ⋅ *s*_*t*−2_ + β_2_ ⋅ *s*_*t*−1_ + β_3_ ⋅ *e*_*t*−1_ + ε and inspected the β_1_ coefficient. Across participants, the mean ± SD of β_1_ was 0.12 ± 0.26 but only significantly different from 0 for one participant. As a further check, we assumed that there was an effect of *s*_*t*−2_ (i.e., an edge between *s*_*t*−2_ and *e*_*t*_ in the causal graph) and added this variable as a regressor to the multiple linear regression model such that *e*_*t*_ and *s*_*t*−1_ were *d*-separated. We found similar mean coefficients for the central tendency and serial dependence effects. As adding this regressor would introduce more multicollinearity in the regression model, we decided to not include the regressor in the final model. The high amount of autocorrelation in the stimulus sequence comes with the disadvantage of a reduced effective variance in the stimulus distances (see Figs 5 and 7) and therefore a reduced precision in the estimated regression coefficients. A possible solution could be to test an experimental condition with a medium amount of autocorrelation.

In conclusion, our findings indicate that the reproduced distances in the vestibular path integration task generally showed central tendency and attractive serial dependence. These perceptual biases were not affected by the level of stimulus autocorrelation, given that covariation of these biases through the stimulus autocorrelation as well as other covariates were taken into account in the model. This suggests that central tendency and serial dependence in vestibular path integration have a neurocognitive rather than a statistical origin.

## Appendix

To compare different central tendency and serial dependence metrics, we simulated reproductions that show central tendency but no serial dependence, i.e., reproductions that tend towards the mean of the underlying stimulus distribution but that are independent of the stimulus distance presented on the previous trial. Such reproductions can be generated for a trial *t* using the following ‘static’ Bayesian model [32]:

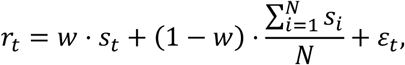

where *r* refers to the reproduced distance, *s* to the stimulus distance, *N* to the total number of trials and ε to a small amount of normally distributed random noise centered on 0. Parameter *w* reflects the weighting between the stimulus on the current trial and the constant mean of all stimuli. The amount of central tendency in the reproductions is defined as *c* = 1 – *w* and the serial dependence is always 0 as the current reproduction does not depend on the previous stimulus irrespective of the amount of central tendency. One simulation for a given *w* consisted of generating reproductions with the static model for a random-walk sequence of 130 stimulus distances (the high-autocorrelation condition) and then shuffling the resulting stimulus-reproduction pairs to create the no-autocorrelation condition. Next, the amount of central tendency and serial dependence in the simulated reproductions was computed using two different methods. First, we used two separate linear least-squares regressions. Central tendency was defined as the slope of the linear regression of the reproduction error (reproduced - stimulus distance) on the stimulus distance. Serial dependence was defined as the slope of the linear regression of the reproduction error of the current trial on the stimulus distance of the previous trial. Second, we computed central tendency and serial dependence as the partial regression coefficients *CT* and *SD_1_* in the multiple linear regression model described in the Methods. We performed 1000 simulations for *w* = 0, *w* = 0.5 and *w* = 1, and we report the mean of the central tendency and serial dependence values across simulations for both methods.

The results are presented in Table A1. Both the simple and multiple linear regression methods compute the correct amount of central tendency (i.e., *c*) in both conditions. Note that the central tendency values are negative as central tendency is defined in terms of the effect of the stimulus distance on the reproduction error (see Fig 1A). Both methods also result in the correct amount of serial dependence (i.e., 0) in the no-autocorrelation condition (see Fig A1, A-C). However, in the high-autocorrelation condition, the central tendency in the reproductions manifests as repulsive serial dependence when computed using the simple linear regression method. This is illustrated in Fig A1, D-F, for three example simulations with increasing amounts of central tendency. If we instead compute serial dependence using the multiple linear regression method in which we control for the current stimulus as well as other variables (see Methods), the resulting value is on average close to 0 across simulations (see Table A1).

**Table A1.**
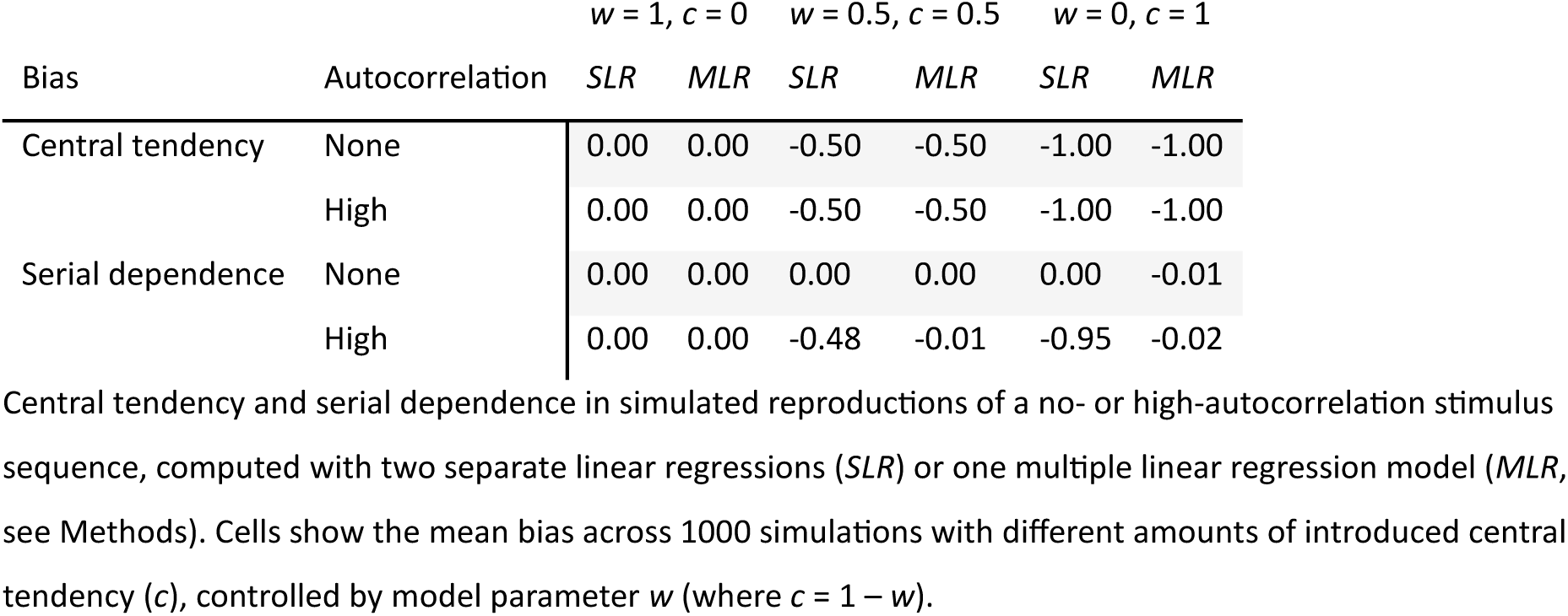
Central tendency and serial dependence coefficients in simulated reproductions.

**Fig A1.**
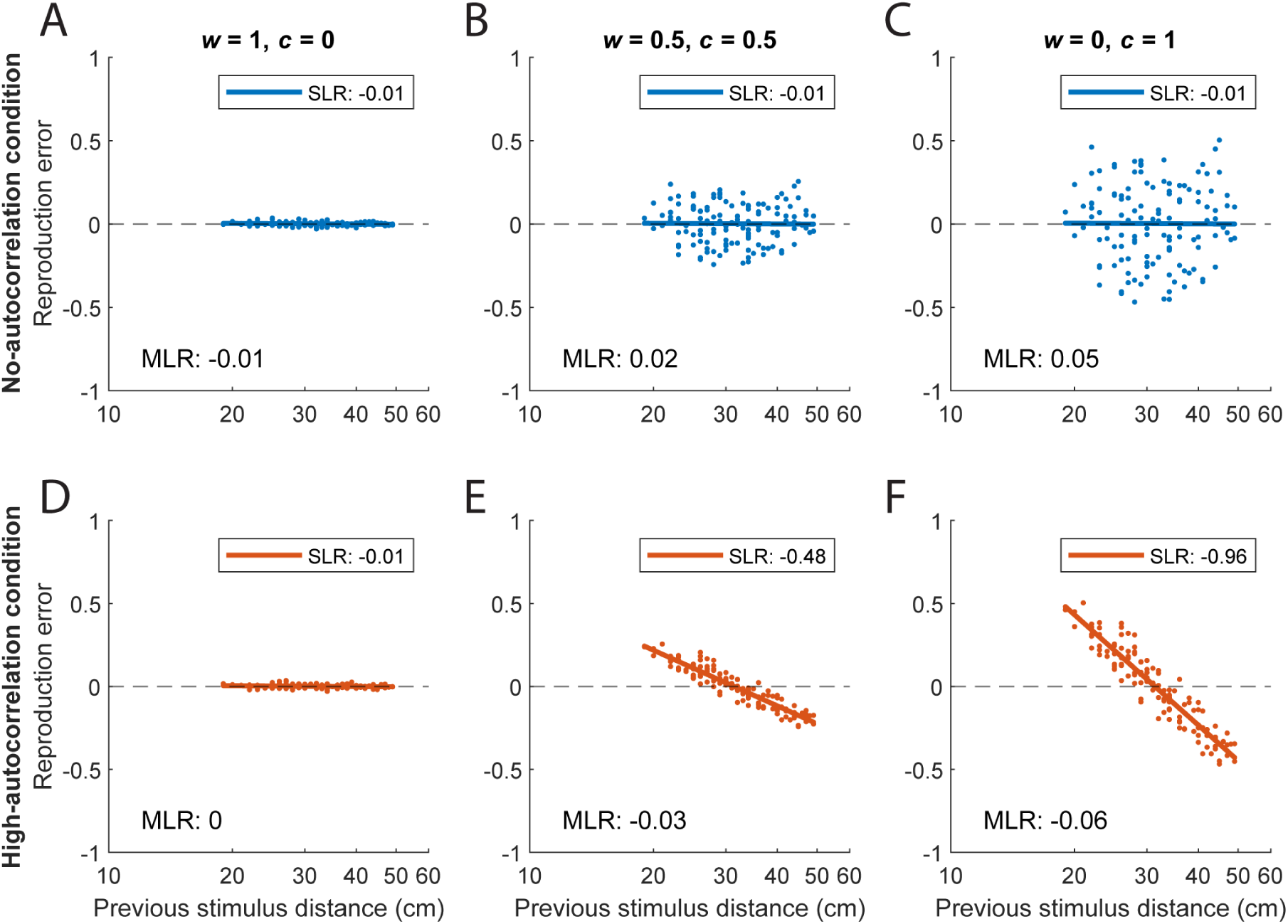
Serial dependence in simulated reproductions. The serial dependence in the no-(blue) and high-autocorrelation (orange) conditions of three example simulations with different amounts of introduced central tendency (*c*), controlled by model parameter *w* (where *c* = 1 – *w*). Serial dependence is plotted as reproduction error (*e*_*t*_ = *r*_*t*_ − *s*_*t*_) against previous stimulus distance *s*_*t*−1_ on logarithmic scale. The slope of the simple linear regression (*SLR*) between these two variables is reported in the key. The corresponding serial dependence value as computed with the multiple linear regression (*MLR*) method (the partial regression coefficient *SD1*, see Methods) is reported in each panel (but not plotted as this coefficient can only be correctly shown in a partial regression plot, see Methods).

## Acknowledgements

We would like to thank Aslan Bellmann for the insightful discussions on causal and statistical testing.

## Author contributions

SCMJW, LOW, RJvB, MK, and WPM conceived and designed research; SCMJW performed experiments; SCMJW analyzed data; SCMJW, LOW, RJvB, MK, and WPM interpreted results of experiments; SCMJW prepared figures; SCMJW drafted manuscript; SCMJW, LOW, RJvB, MK, and WPM edited and revised manuscript; SCMJW, LOW, RJvB, MK, and WPM approved final version of manuscript.

## Data availability statement

All data and code are available for review in the Radboud Data Repository collection: di.dcc.DSC_2024.00135_205. Upon publication, all data and code will be made publicly available via the DOI reserved for this collection: https://doi.org/10.34973/w21g-wt67.

